# Structural basis of mammalian Complex IV inhibition by steroids

**DOI:** 10.1101/2022.04.13.488242

**Authors:** Justin Di Trani, Agnes Moe, Daniel Riepl, Patricia Saura, Ville R. I. Kaila, Peter Brzezinski, John L. Rubinstein

## Abstract

The mitochondrial electron transport chain maintains the proton motive force that powers ATP synthesis. The energy for this process comes from oxidation of NADH and succinate, with the electrons from this oxidation passed via intermediate carriers to oxygen. Complex IV (CIV), the terminal oxidase, transfers electrons from the intermediate electron carrier cytochrome *c* to oxygen, contributing to the proton motive force in the process. Within CIV, protons move through the K- and D-pathways during turnover. The former is responsible for transferring two protons to the enzyme’s catalytic site upon reduction of the site, where they eventually combine with oxygen and electrons to form water. CIV is the main site for respiratory regulation, and although previous studies showed that steroid-binding can regulate CIV activity little is known about how this regulation occurs. Here we characterize the interaction between CIV and steroids using a combination of kinetic experiments, structure determination, and molecular simulations. We show that molecules with a sterol moiety, such as glyco-diosgenin and cholesteryl hemisuccinate, reversibly inhibit CIV. Flash photolysis experiments probing the high-speed equilibration of electrons within CIV demonstrate that binding of these molecules inhibits proton uptake through the K-pathway. Single particle cryo-EM of CIV with glyco-diosgenin reveals a previously undescribed steroid-binding site adjacent to the K-pathway, and molecular simulations suggest that the steroid binding modulates the conformational dynamics of key residues and proton transfer kinetics within this pathway. The binding pose of the sterol group sheds light on possible structural gating mechanisms in the CIV catalytic cycle.

**SIGNIFICANCE STATEMENT:** Mammalian complex IV (CIV), the final complex of the mitochondrial electron transport chain, uses electrons from cytochrome *c* to reduce oxygen to water, driving aerobic life. Although CIV functions as the main site for respiratory regulation, there is little structural or biochemical information on how this regulation occurs. Previous studies provided evidence of CIV regulation by steroids, but the steroid binding site and regulatory mechanism remain unclear. Using single particle cryogenic electron microscopy, we discover the binding site of the steroid-derived detergent, glyco-diosgenin. Results from flash photolysis kinetic experiments with CIV in the presence of glyco-diosgenin and cholesterol hemisuccinate are combined with cryo-EM and molecular simulations to elucidate how steroid binding limits proton uptake by the complex.

## INTRODUCTION

The transmembrane protein complexes of the electron transport chain power the production of ATP by maintaining an electrochemical proton motive force (PMF) across the mitochondrial inner membrane or the plasma membrane of bacteria. The first component of the mitochondrial chain, complex I, initiates electron transport by oxidizing NADH produced by the Krebs cycle, glycolysis, and fatty acid oxidation. Complex I uses electrons from NADH to reduce membrane bound quinone to quinol while pumping protons from the negatively charged matrix side (N side) of the mitochondrial inner membrane to the positively charged intermembrane space side (P side). Complex II oxidizes succinate to fumarate and reduces quinone, contributing to the quinol pool. Complex III oxidizes quinol to quinone through a mechanism known as the Q cycle, adding to the PMF and reducing soluble cytochrome *c* in the intermembrane space. Reduced cytochrome *c* then binds to complex IV (CIV), the terminal enzyme of the electron transport chain. CIV contributes to the PMF while transferring electrons from cytochrome *c* to the final electron acceptor, O_2_, which is reduced to H_2_O. Each electron transfer from cytochrome *c* to O_2_ is linked to proton uptake from the N side and pumping of one proton from the N to the P side of the membrane. Thus, overall, the eukaryotic electron-transport chain couples proton translocation across the mitochondrial inner membrane with electron transfer from NADH and succinate to O_2_.

Reduced cytochrome *c* binds to CIV and is oxidized by its primary electron acceptor, Cu_A_. The electron is then transferred to heme *a* and onwards to the enzyme’s catalytic site, which includes heme *a*_3_ and Cu_B_ (**Fig 1A**, *top left*)(for reviews see (1, 2)). Overall, CIV catalyzes the oxidation of four cytochrome *c* molecules, uptake of eight protons from the N side, and reduction of one dioxygen molecule to produce two water molecules and add four protons to the P side. Turnover of CIV can be divided into a reductive phase and an oxidative phase. In the reductive phase, two electrons are transferred from cytochrome *c* molecules to the oxidized catalytic site (**Fig. 1A**, *top right*). In the oxidative phase, dioxygen binds to the reduced heme *a*_3_ and CIV receives two more electrons from cytochrome *c* molecules to complete the reaction cycle (**Fig. 1A**, *bottom*). Reduction of the oxidized catalytic site is coupled to the transfer of two protons from the N side to the catalytic site through the K-pathway (3, 4), which comprises conserved residues including the Lys319 residue (*B. taurus* CIV amino-acid residue numbering) that gives the pathway its name (5). After binding of dioxygen to the catalytic site and transfer of the next two electrons from cytochrome *c* molecules, proton transfer from the N side to the catalytic site occurs through the D-pathway. The D-pathway is composed of water molecules and conserved polar and protonatable residues, including Glu242 and Asp91, the latter giving the pathway its name (6). During each electron transfer from cytochrome *c* to the catalytic site, protons are pumped across the membrane through the D-pathway (for review, see (2, 7–11)). A third pathway (the H-pathway) has also been suggested for the mammalian CIV (8, 12), but remains controversial because it is not present in the mitochondrial CIV from *Saccharomyces cerevisiae* (13) or bacterial CIVs (14).

**Fig. 1.**
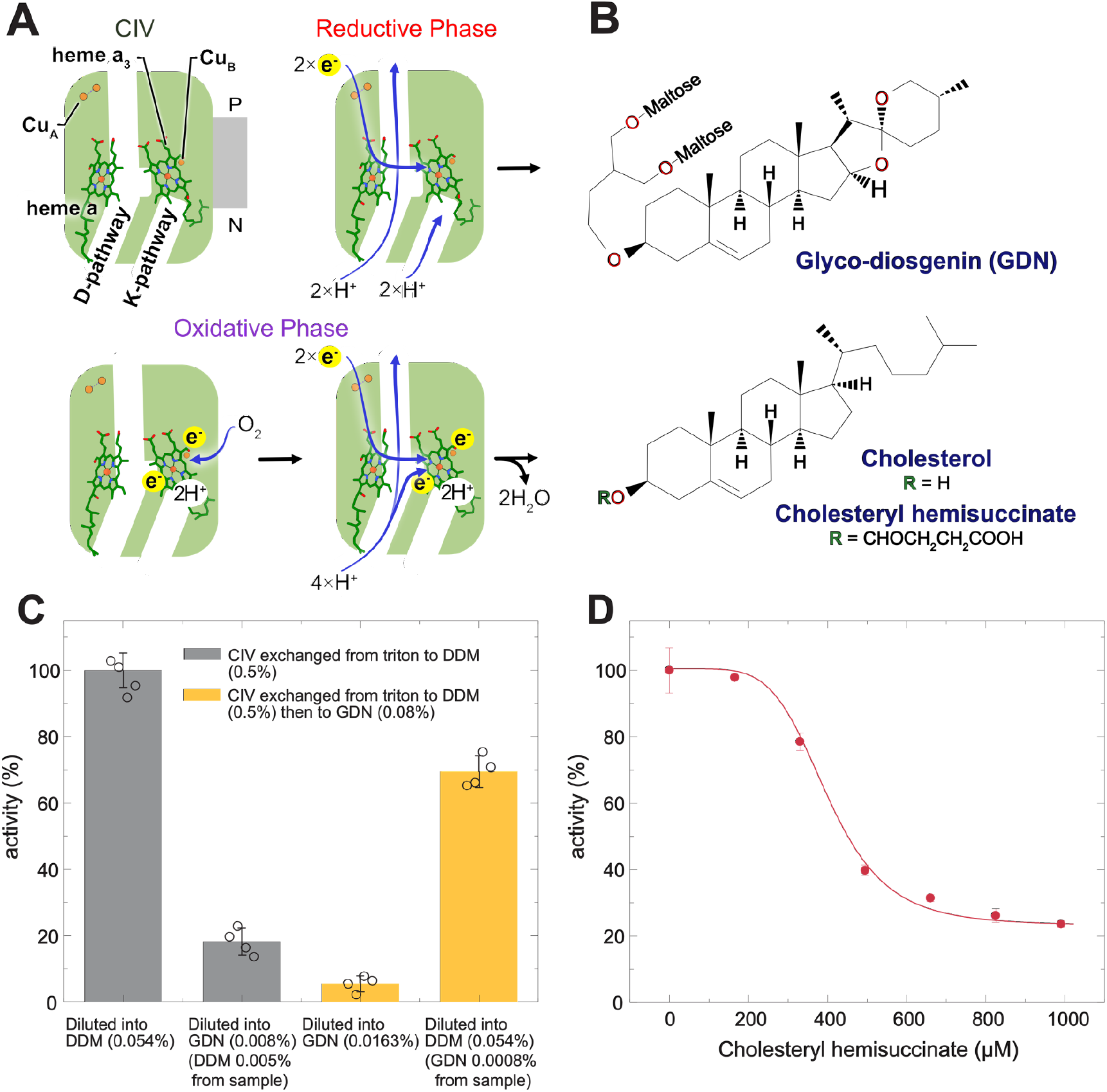
Inhibition of CIV by cholesterol analogues. **A**, Cartoon of CIV structure, with electron carriers and channels labelled (*top left structure*). In the reductive phase, CIV pumps protons across the inner membrane through the D-pathway and transfers protons to the catalytic site through the K-pathway (*top right structure*). In the oxidative phase, oxygen binds to the catalytic site (*bottom right structure*) followed by transfer of protons across the inner membrane and to the catalytic site through the D-pathway (*bottom right structure*). **B**, Structure of GDN, cholesterol, and cholesteryl hemisuccinate (CHS). **C**, Cytochrome *c* oxidase activity measured with a Clark-type oxygen electrode. CIV incubated in DDM then diluted into DDM (*first grey bar*) or GDN (*second grey bar*), incubated in GDN and diluted into DDM (*first orange bar*) or GDN (*second orange bar*). **D**, CIV activity in LMNG with varying concentrations of CHS (*red points*, mean ± s.d., n=3 independent measurements), and fit to data (*red line*).

Complex IV is thought to be the central regulatory site for oxidative phosphorylation in mitochondria (15). This regulation is of paramount importance as mitochondrial dysfunction can lead to the progression of numerous human diseases including neuro-degeneration, ischemia reperfusion injury, cancer, and diabetes (15). Mammalian CIV is regulated by phosphorylation of subunits (16) and differential expression of subunit isoforms (17). CIV also possesses conserved lipid binding sites (18) that allow the enzyme’s lipid environment to modulate its activity (19, 20). Lipophilic small molecules, including detergents, steroids, and bile salts, have a complex effect on the activity of CIV from *Rhodobacter sphaeroides* (18, 21). Steroids as well as thyroid hormones have been shown to inhibit mammalian CIV (22, 23), possibly by interfering with K-pathway proton translocation (24), thyroid hormones also appear to suppress superoxide formation by CIV (23). This effect has also been seen when mitochondria are exposed to steroids and glucocorticoids, which causes decreased oxygen consumption, an effect that is being explored in relation to cancer therapies (25, 26). Further, accumulation of bilirubin inhibits CIV, possibly by blocking K-pathway proton translocation, which was suggested to be linked with development of neonatal encephalopathy. This inhibition can be reversed by the bile salt glycoursodeoxycholic acid (10, 27).

Here, we explore the interaction between mammalian CIV and the steroids cholesteryl hemisuccinate (CHS) and glycol-diosgenin (GDN). CIV activity measurements show that both GDN, a detergent with a steroid-based lipophilic group (28), as well as CHS, reversibly inhibit CIV. Flash photolysis reveals that binding of each of these compounds results in impaired K-pathway proton translocation, similar to K-pathway mutants in *R. sphaeroides*. The cryo-EM structure of CIV with GDN defines the sterol binding site and pose, which is different from a site previously proposed from crystal structures of proteins purified and crystallized with sodium cholate and deoxycholate (29–32). Importantly, the binding site is adjacent to helix VII of subunit I, which is in close contact with Lys319 and may have a direct role in proton gating. Molecular simulations explain the inhibitory effect of sterol binding on proton transfer through the K-pathway at a molecular level.

## RESULTS AND DISCUSSION

### Inhibition of mammalian CIV by cholesterol derivatives

A number of bile acids and other sterol-containing compounds, including detergents, have been shown to inhibit mammalian and bacterial CIV activity *in vitro* (21, 22). To explore this effect, CIV was purified from bovine heart mitochondria (33) and its activity was assayed with a Clark electrode, which measures the reduction of oxygen as a function of time. Concentrated CIV was diluted into an aerobic buffer containing a pool of cytochrome *c*, which is kept reduced by an excess of ascorbate and the electron mediator *N,N,N*’,*N*’-tetramethyl-*p*-phenylenediamine (TMPD). Under these conditions, CIV uses the electrons from cytochrome *c* to reduce dissolved oxygen until it is depleted. Clark electrode measurements with CIV purified in triton X-100 and exchanged into *n*-Dodecyl-B-D-maltoside (DDM) showed an electron transfer rate of 220 ± 10 e^-^/s (mean ± s.d., *n*=4 measurements from two separate purifications), similar to previous studies in DDM (33). However, upon dilution of concentrated DDM-solubilized CIV into buffer containing glyco-diosgenin (GDN), which includes a steroid-derived lipophilic group (**Fig. 1B**, *top*)(28), CIV activity decreased by ∼80% (**Fig. 1C**, *second bar*). CIV exchanged into GDN buffer during ion exchange chromatography and assayed in buffer containing GDN displays only ∼5% of the activity of the DDM-solubilized CIV assayed in DDM-containing buffer (**Fig. 1C**, *third bar*), but upon dilution into DDM buffer it regains ∼60% of the activity of DDM-solubilized CIV assayed in DDM-containing buffer (**Fig. 1C**, *fourth bar*). These data show that GDN inhibits CIV, and that DDM can reverse the inhibition. This observation is consistent with previous data showing reversible inhibition of mammalian CIV by sterol-containing compounds (22). The ability of DDM to reverse inhibition of CIV by small molecules was demonstrated with CIV from *R. sphaeroides* (21, 34), and is likely related to DDM’s ability to strip hydrophobic molecules from proteins (35).

To confirm that the observed inhibition was caused by the steroid moiety of GDN, CIV was titrated with cholesteryl hemisuccinate (CHS), a cholesterol derivative that is more soluble than cholesterol due to the addition of a succinate group (**Fig. 1B**, *bottom*). CIV in DDM was exchanged to the lower critical micelle concentration (CMC) detergent, lauryl maltose neopentyl glycol (LMNG), before titration with CHS. Detergents with lower CMCs are often considered more ‘mild’ than higher CMC detergents and exchange from DDM to LMNG was performed to minimize the risk of removal of CHS from CIV by the detergent. In the absence of CHS, the activity of CIV in LMNG was 200 ± 15 e^-^/s (mean ± s.d., *n*=4 measurements from two separate purifications), comparable to its activity in DDM. As CHS was gradually increased to 990 μM, the activity of CIV decreased to ∼20% of its initial activity (45 ± 5 e^-^/s) (**Fig. 1D**). Together, these data show that the sterol-containing molecules GDN and CHS can also inhibit CIV, likely due to the sterol moieties of the molecules.

### Steroids inhibit proton uptake through the K-pathway

The absorbance spectra of the redox cofactors in CIV change with redox state, making it possible to determine how electrons are distributed amongst the internal electron carriers. Stopped-flow results from earlier measurements with *R. sphaeroides* CIV revealed that bile salts caused accumulation of electrons at heme *a*, suggesting that the electron transfer from heme *a* to the catalytic site is rate-limiting (21, 23, 36).

To identify the internal electron transfers that are inhibited by GDN and CHS, we investigated intramolecular electron transfer and proton-coupled electron transfer reactions in CIV with the so-called flow-flash technique (see e.g. (4)). To study the oxidative phase with this technique, CIV is reduced in the absence of O_2_ and the catalytic site is blocked by binding of carbon monoxide (CO). After addition of O_2_, the CO ligand is removed by a short laser flash. Removal of CO allows O_2_ to bind to the catalytic site and initiate oxidation of the enzyme, which can be followed spectrophotometrically with μs time resolution. As indicated above, in the oxidative phase all protons are transferred through the D-pathway and any effects from slowed proton uptake are typically manifested in slowing electron transfers that are linked to D-pathway proton uptake (37). The time constant of these components (***SI Appendix* Fig. S1**) were the same in GDN and DDM, indicating that the impaired activity of CIV in GDN is not due to slowed intramolecular electron transfer or proton uptake through the D-pathway.

The reductive phase is typically investigated by mixing oxidized CIV with reduced cytochrome *c*. However, in this reaction, electron transfer from cytochrome *c* to CIV is rate limiting, which precludes identification of specific intramolecular reactions that may be slowed. To overcome this limitation, we measured internal electron transfer and proton-coupled electron transfer in flash photolysis experiments (38, 39). In these experiments, CIV is incubated with CO in an anaerobic environment creating a mixed valence state where Cu_A_ and heme *a* are oxidized, while Cu_B_ and heme *a*_3_ at the catalytic site are reduced, and CO is bound to heme *a*_3_ (**Fig. 2A**, *top left*)(4). The heme *a*_3_-CO bond is photolabile and upon a short laser flash the CO ligand dissociates and equilibrates with the gas dissolved in solution (**Fig. 2A**, *pink arrow*). Upon dissociation of CO, the apparent midpoint potential of heme *a*_3_ decreases, which results in electron transfer from heme *a*_3_ to heme *a* and then to Cu_A_ over time scales of ∼3 μs and 30 μs, respectively (4). Because this electron transfer occurs in the opposite direction of the electron transfer during turnover, the reaction is referred to as “electron backflow”. However, the rate constants of the kinetic components reflect the sum of the forward and backward rate constants and therefore provide information on intramolecular electron transfer in the reductive phase. With the mammalian CIV, after CO dissociation the difference in heme *a*/*a*_3_ midpoint potentials yields ∼10-30% (depending on the source of CIV) oxidation of heme *a*_3_ (38, 40), with the remaining population retaining the electron at heme *a*_3_ (**Fig. 2A**, *bottom*). In this remaining population, the electron equilibrium shifts further toward heme *a* once a proton is transferred through the K-pathway (4, 38) (**Fig. 2A**, *purple arrow*). Both the fraction of this component and its time constant increase with increasing pH (p*K*_a_ ≅ 8-9, depending on the source of CIV) (38, 40), and therefore the reaction is monitored at pH>7. This electron transfer event can be monitored by measuring the change in absorbance at 598 nm (38).

**Fig. 2.**
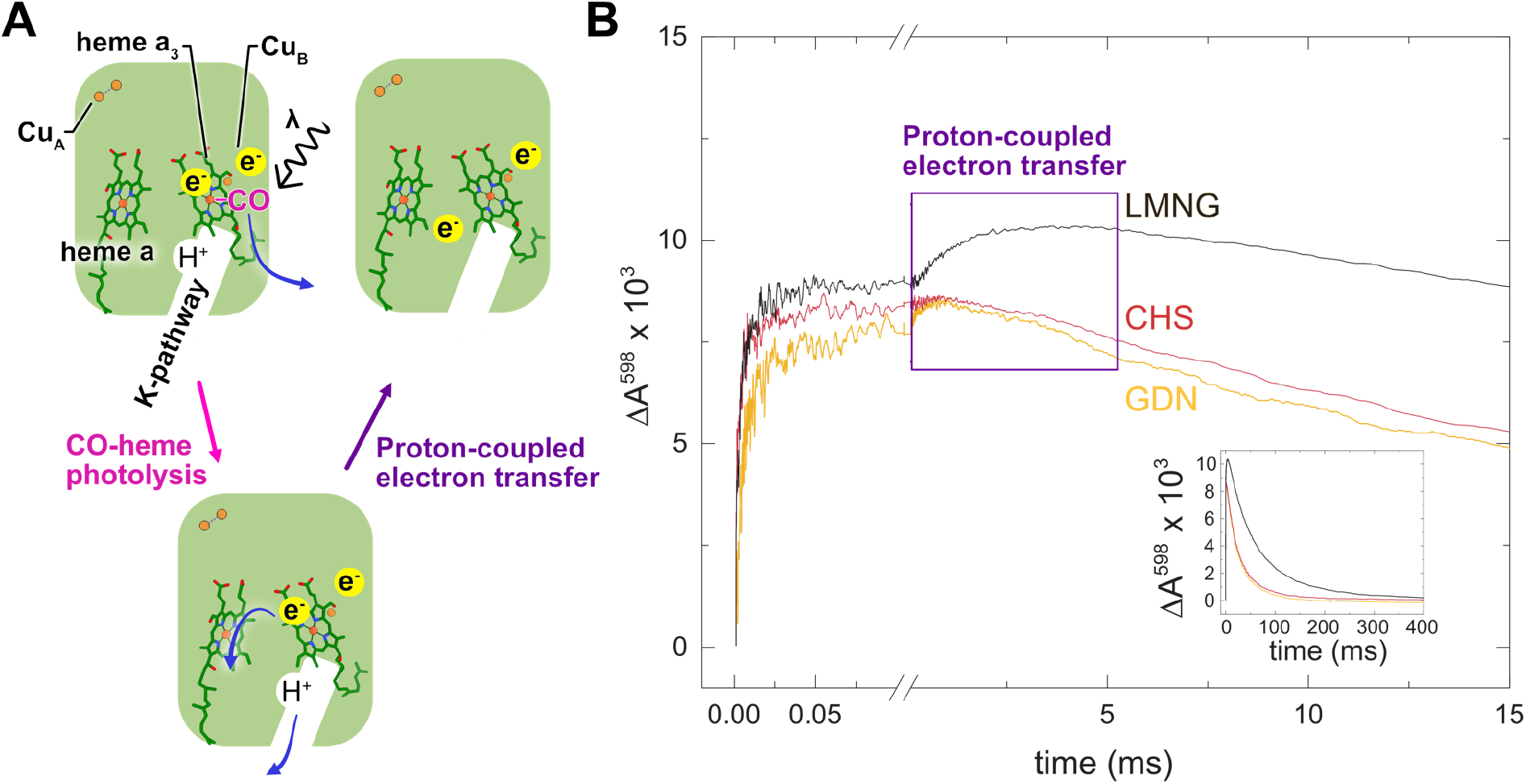
Flash photolysis experiments. **A**, CIV in the mixed valence state (*top left structure*) is excited by laser flash (*pink arrow*), in a population where heme *a*_3_ remains reduced electron transfer from heme *a*_3_ to heme *a* is coupled with proton translocation through the K-pathway (*purple arrow*), producing a change in absorbance at 598 nm. **B**, Absorbance changes at 598 nm. The data within the purple box show a kinetic phase attributed to proton-coupled electron transfer from heme *a*_3_ to heme *a* for LMNG only (*black curve*), LMNG + CHS (*red curve*), and GDN (orange curve). Inset shows zoom out of absorbance changes over 400 ms.

To investigate the effect of sterol binding on the reductive phase, CIV in LMNG was first prepared in the mixed-valence state for flash photolysis experiments. Upon laser-induced photolysis, CO disassociates and an electron is transferred from heme *a*_3_ to heme *a* in a subpopulation of CIV complexes, causing a large increase in the absorbance at 598 nm in the time range 0-50 μs (38) (**Fig. 2B**, *black curve, first 0*.*05 ms*). In the remaining subpopulation of CIV complexes a proton must leave through the K-pathway for the electron transfer to occur with a time constant of 1.6 ± 0.2 ms (average of ∼200 traces from each of two separate purifications) (**Fig. 2A**, *purple arrow*, and **Fig. 2B**, *black curve in purple box*). Over a longer time scale the signal at 598 nm slowly decreases back to baseline as CO recombines with CIV (**Fig. 2B**, *black curve, inset*) bringing the system to equilibrium with a time constant of ∼58 ms and allowing for repetition of the experiment. CIV with LMNG and CHS, as well as CIV exchanged to GDN, were also prepared in the mixed valence state. Upon flash photolysis, both samples showed a rapid increase in absorbance in the 0-50 μs time range, indicating that the photolysis and electron transfer occurs in a portion of the population. This observation shows that internal electron transfer is not affected by binding of GDN or CHS (**Fig. 2B**, *blue and red curves, first 50 μs*). However, after this rapid increase in absorbance there is no further absorbance increase, so that the 1.6-ms component is not observed (**Fig. 2B**, *blue and red curves in purple box*). The absence of this component demonstrates that the proton-coupled electron transfer does not occur. These results show that proton transfer through the K-pathway is blocked in the presence of GDN or CHS. Furthermore, this loss of the 1.6-ms component is identical to data from flash photolysis studies in which the K-pathway of CIV from *R. sphaeroides* was deactivated by site-directed mutagenesis (4). Therefore, these data also support the suggestion that slowed proton uptake through the K-pathway is the reason for accumulation of electrons at heme *a* when *R. sphaeroides* CIV is treated with bile salts (21, 36). As shown in the inset in **Fig. 2B**, recombination of the CO with the LMNG sample was a factor of ∼2 (∼58 ms) slower than with the LMNG+CHS and CHS samples (∼28 ms). As shown previously (38), the CO ligand binds only to the fraction of CIV in which heme *a*_3_ is reduced, yielding a CO-recombination time constant that is dependent on the heme *a*_3_/heme *a* electron equilibrium. Because GDN and CHS block proton-coupled electron transfer from heme *a*_3_ to heme *a*, the fraction heme *a*_3_ that is reduced increases and CO recombination with CO can occur faster.

### Structure of GDN inhibited CIV reveals steroid proximity to K-pathway

To gain insight into how steroids limit proton translocation along the K-pathway, CIV in GDN was subjected to structural analysis by cryo-EM. GDN was selected over LMNG with CHS because of the possibility that LMNG could compete with CHS for interaction with the sterol-binding site, while with CIV in GDN there is no possibility for this competition. To detect conformational changes induced by reduction of the complex not seen in crystal structures, CIV was reduced and inhibited with CO prior to freezing cryo-EM specimens. Subsequent imaging and image analysis led to a 3.1 Å resolution map of the complex (**Fig. 3A**, *top*, ***SI appendix* Fig. S2A-D** and **Table S1**). The map of CIV showed density for three core and ten peripheral subunits of the complex, but is missing the NDUFA4 subunit originally thought to be part of complex I (41). The map fit previous atomic models of CIV well (**Fig. 3A, *SI Appendix* Fig. S2E**) and no conformational differences were apparent between the present structure and earlier atomic models of CIV (42). In addition, 3D variability analysis (43) did not detect any motions, suggesting a rigid protein structure. Clear density was apparent for the lipids phosphatidylethanolamine (PE) and phosphatidylglycerol (PG) between subunit I and III (**Fig. 3A**, *yellow models*, ***SI Appendix* Fig. S2E**), which were seen in earlier crystal structures (18, 42). These lipids have been suggested to be important for CIV function (31).

**Fig. 3.**
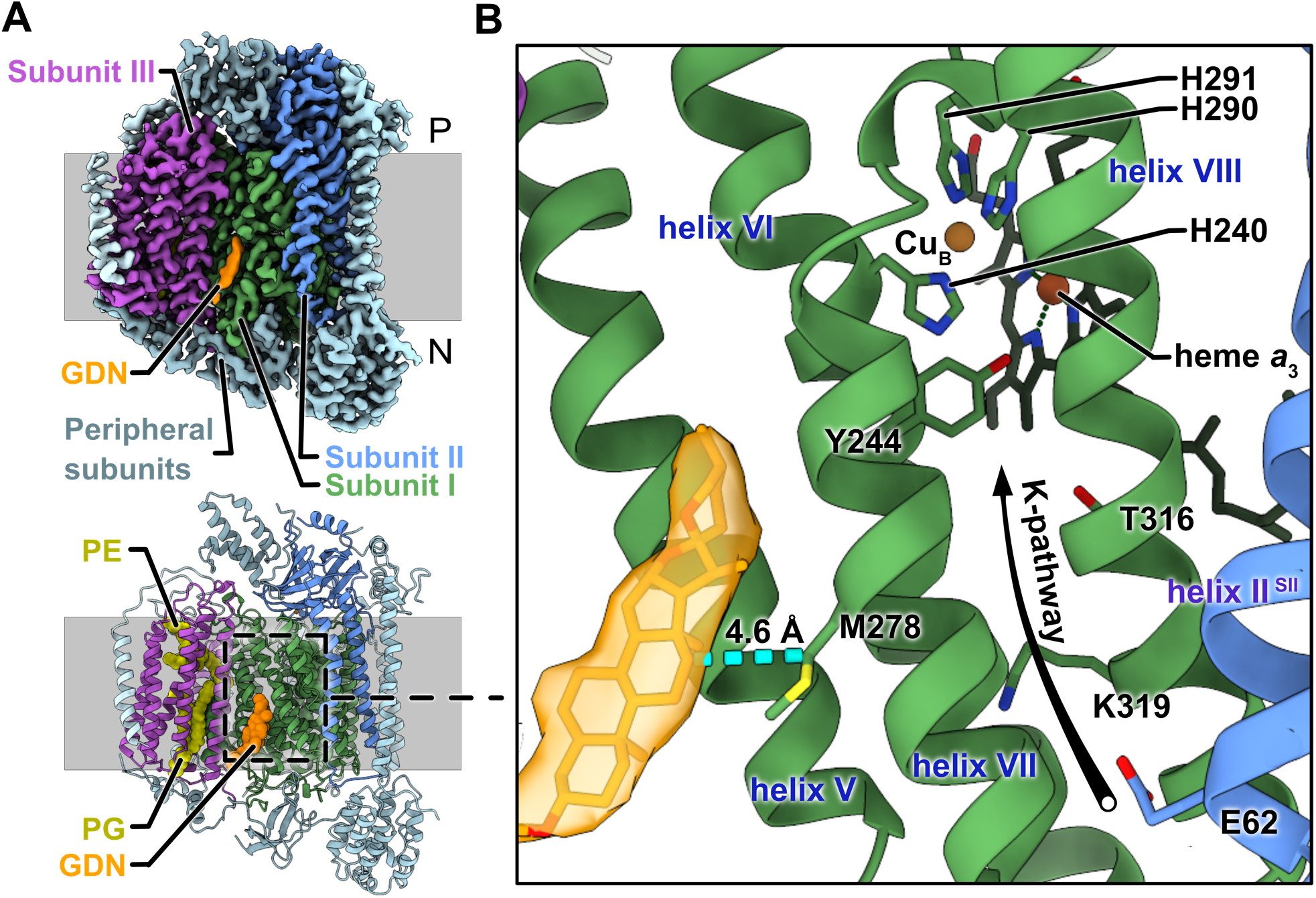
Cryo-EM structure of GDN-inhibited CIV. **A**, Cryo-EM map (top), and atomic model for CIV (*bottom*) with lipids phosphatidylethanolamine (PE) and phosphatidylglycerol (PG) modelled and GDN shown in orange. **B**, Close up of K-pathway with map and model for the GDN (*orange map and model*).

The proton translocation pathways in CIV are formed by the three core subunits (subunit I-III) of CIV. The entrance to the K-pathway is found on the N side of the inner membrane at the interface of helix VII from subunit I and helix II of subunit II (**Fig. 3B**), the latter bearing Glu62, the first amino acid in the K-pathway proton transfer chain. This region of the map is at a lower resolution than surrounding regions (***SI Appendix* Fig. S2D**), consistent with the higher B-factors for residues at the entrance of the K-pathway in earlier crystal structures (21). These higher B-factors, which correspond to uncertainty in the positions of atoms, have been attributed to movements at the K-pathway opening that are necessary for proton uptake (24). The remainder of the pathway is formed by helices VI and VII of subunit I, bearing the Cu_B_ ligands His240 (helix VI), His290 (helix VII), and His291 (helix VII), as well as helix VIII of subunit I, which contains the eponymous Lys319 residue. Subunit I also contains Thr316 and Tyr244, the latter being part of the catalytic site, which participates in the reduction of oxygen to water (1). Helix VII lies between the K-pathway and the outside of the protein and is in close contact with Lys319 (**Fig. 3B**). The cryo-EM map reveals clear density for the sterol portion of a GDN molecule adjacent to helix VII (**Fig. 3A** and **B**). With the standard deviation of the cryo-EM map normalized to σ = 1, the sterol is apparent at 8σ, similar to the density from helix VII, suggesting high occupancy of GDN in this site. However, no clear density corresponding to the maltose groups of GDN could be detected, suggesting that these portions of the detergent molecule are more dynamic. The sterol from GDN sits in a hydrophobic cleft formed by helices V, VI, and VII of subunit I. This position puts GDN in close contact (∼4.6 Å) with Met278 on helix VII (**Fig. 3B**, *cyan dashed line*) where it is positioned to affect dynamics of the helix and proton translocation as described below.

Cryo-EM shows that the CIV preparation used in the present study was monomeric. Both dimers and monomers of CIV have been observed in solution previously (44–47), with monomers showing more activity than dimers (46). Bile salts such as sodium cholate, which include a sterol moiety, promote formation of CIV dimers (47), with both oligomeric states having been resolved by X-ray crystallography (12, 48). X-ray crystallography of bovine CIV dimers in the presence of cholate showed cholate in the crystals at the dimer interface (21, 49). Diffraction analysis of *R. sphaeroides* CIV crystals grown in the presence of cholate showed cholate between CIV complexes at crystal contacts in the lattice (32). These studies led to the proposal that the dimer interface forms a sterol binding site near the entrance to the K-pathway for both species. More recently, cholate was modelled far from the K-pathway in a X-ray crystal structure of monomeric bovine CIV, indicating that cholate remains in the sample after purification with cholate but is not bound at the dimerization interface (48). Further, the CIV NDUFA4 subunit seen in a recent cryo-EM structure of human CIV (41) is bound where the monomer-monomer interface forms in dimers, blocking the proposed cholate bind site. Together, the inconsistent location of cholate in X-ray crystal structures suggests that the presence of cholate in CIV crystals is due to the crystallization process and does not indicate a specific sterol binding site. The present results show that sterols can indeed bind monomeric CIV. Dimers of bovine CIV appear to exclude steroids from the binding site seen here because the N-terminal tail of subunit VIA in each monomer occupies the sterol-binding pocket of the opposite monomer (***SI Appendix*, Fig. S2F**) (42), which may also explain the lower activity of the dimer (46). In the simpler four subunit CIV from *R. sphaeroides*, the sterol-binding site seen here with bovine CIV monomers is blocked by subunit IV (50), which is not present in the mammalian enzyme.

### Role of helix VII in K-pathway gating

Proton transfer along the K-and D-pathways requires continuous hydrogen-bonded chains of water molecules as well as polar and protonatable amino acid side chains. While no water molecules could be resolved in the current map, crystal structures of CIV show that the D-pathway is highly hydrated (51). The K-pathway, on the other hand, shows gaps in the water networks (42, 51, 52). This lack of a continuous proton-conducting pathway implies that reorientation of sidechains and hydration must occur to accommodate proton transfer through the K-pathway (51, 53, 54). Prior molecular dynamics (MD) simulations showed that Lys319 can adopt multiple conformations, with the deprotonated side chain adopting a ‘down’ conformation far from Tyr244 and the protonated side chain adopting an ‘up’ conformation near to Tyr244, which allows for water-mediated proton transfer to the catalytic site (54– 56).

To understand how GDN binding affects proton transfer through the K-pathway, we performed MD simulations of the bovine CIV with and without GDN modeled next to helices VI and VII in the orientation observed by cryo-EM (**Fig. 4A**). Similar to previous simulations, Lys319 adopts both an up and down conformation (**Fig. 4A**). During the MD simulations without GDN, up to ∼10 water molecules diffuse into the K-pathway from the N side of the membrane (**Fig. 4B, *SI Appendix* Fig. S3A**, *simulation 5*). The Lys319 sidechain flips from its down conformation (Lys319-Tyr244 distance > 16 Å) to its up conformation (Lys319-Tyr244 distance < 12 Å), which results in formation of a hydrogen-bonded water wire with 3-4 water molecules leading from Lys319 to Tyr244 via Thr316 (**Fig. 4B**, *circled in blue*). These 3-4 water molecules allow for proton transfer between Lys319 to Tyr244. Re-orientation of Lys319 into its up conformations is observed only in simulations where the Cu_B_ is reduced (54), whereas with an oxidized catalytic site Lys319 samples just the down conformation far from Y244, consistent with previous results (54, 57, 58). In contrast to the up conformation of Lys319 in the absence of GDN (**Fig. 4C**, *blue curve*), when the MD simulations are performed with GDN modeled between helices VI and VII, Lys319 samples only the down orientation, independent of the modeled redox state at Cu_B_ (**Fig. 4C**, *orange curve*). This finding, in addition to previous simulations (58), are also consistent with recent cryo-EM structures of CIV from *Paracoccus denitrificans* showing that Lys319 moves towards Try244 (280 in *P. denitrificans*) when the enzyme is in the reduced state (59).

**Fig. 4.**
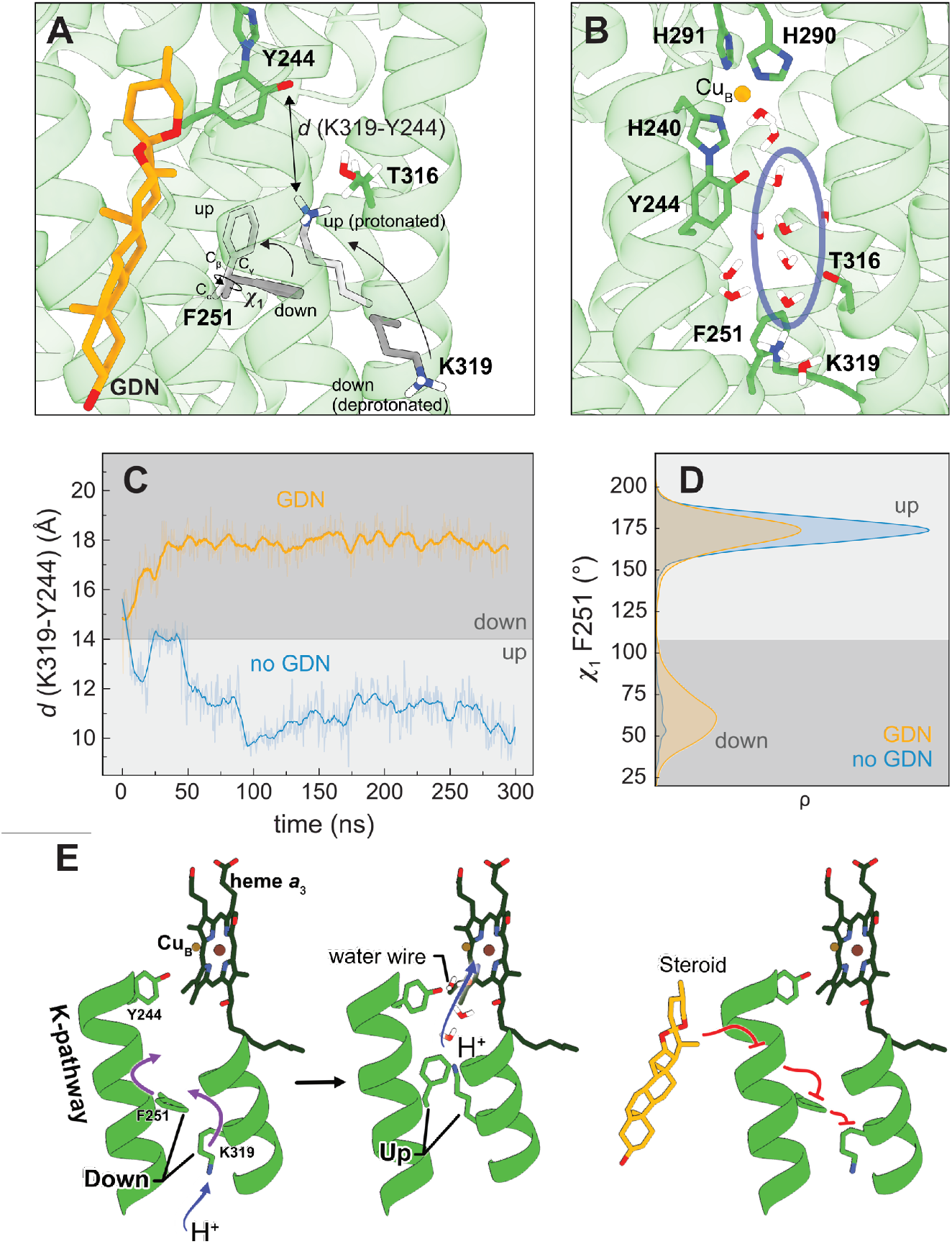
Molecular simulations of CIV K-pathway dynamics and proton uptake. **A**, Snapshot of MD trajectories. Phe251 and Lys319 are displayed in the ‘down’ conformation (*dark grey*) and the ‘up’ conformation (*light grey*), with the Lys319-Y244 distance labelled. **B**, Residues involved in the proton transfer reaction, showing the structure of the water wire connecting Lys319 and Cu_B_, with the water molecules between Lys319 and Tyr244 circled in *blue*. **C**, Simulation trajectories in the presence (*orange curve*) and absence (*blue curve*) of GDN. In the presence of GDN, Lys319 samples the down conformation (*d* > 16 Å), in the absence of GDN, Lys319 can flip to its up conformation (*d* < 12 Å). **D**, Distribution of the Phe251 sidechain dihedral angle (*χ*_1_) from simulations 1-6 (see SI Appendix, Table S2). Simulations with GDN show an enhanced sampling of the down conformation of Phe251 (*χ*_1_=60°) as compared to the simulations without a sterol. The down conformation of Phe251 could prevent Lys319 from flipping upwards (see panel B) and suppresses proton uptake through the K-pathway. **E**, When Phe251 and Lys319 are in the down conformation Lys319 can extract a proton form the N side of the inner membrane (*left structure)*. The correlated movement of Phe251 and Lys319 to the up conformation allows proton transfer to the catalytic site (*middle structure*). Steroids block movement of Phe251, which in turn blocks Lys319, preventing proton transfer through the K-pathway.

The simulation trajectories suggest that GDN binding affects the conformation of aliphatic residues in helices VI and VII, which in turn affect the conformation of Lys319. In particular, Phe251 on helix VI samples a down orientation (ρ_1_ of Phe251 ∼170°) in the presence of GDN (**Fig. 4D**, *orange*, ***SI Appendix* Fig. S3B-D**), which appears to block rotation of Lys319 into its up conformation. In contrast, in the absence of GDN the aromatic sidechain is only found in the up conformation (**Fig. 4D**, *blue*). Simulations with cholesterol in the sterol binding cavity show similar results to those with GDN (***SI Appendix* Fig. S3C**,**D**, *red*). Therefore, the up conformation of Phe251 appears to be required for Lys319 to adopt its up conformation, which in turn is needed for water wire formation and proton translocation (**Fig. 4E**, *left two structures*). GDN binding alters the orientation of Phe251, which in turn blocks Lys319 and prevents proton translocation (**Fig. 4E**, *Right structure*). Together, these results show how GDN binding to the site observed by cryo-EM induces the changes in internal electron transfers seen in kinetic experiments.

## MATERIALS AND METHODS

### Preparation of bovine heart mitochondria

Bovine hearts were kept on ice and all subsequent steps were carried out at 4°C. Fat, blood vessels and connective tissues were removed from the hearts and the remaining material was cut into ∼2 cm^3^ pieces before being ground with a meat grinder. Portions of 200 g of ground heart were mixed with 600 mL of buffer (250 mM sucrose and 10 mM Tris pH 7.8) and 3 mL of 2 M Tris (pH unadjusted) in a blender for 45 s. 4 mL of 2 M Tris was then added and mixed for another 45 s and pH was adjusted to 7.8 if necessary. The homogenate was centrifuged at 1,600 × g for 15 min to remove unbroken cells and nuclei. The supernatant was filtered through two layers of cheesecloth to remove lipid granules and centrifuged at 17,000 × g for 30 min. The pellet was then resuspended in 250 mM sucrose and 10 mM Tris pH 7.8 and centrifuged at 17,000 × g for 30 min. The resulting pellet consisted of two layers, one soft and light in colour, and one darker and firmer. The lighter fraction (broken mitochondrial fragments) was removed by decantation. The procedure was then repeated until only the heavy fraction of mitochondria remained after centrifugation. The heavy fraction of mitochondria was frozen in liquid nitrogen and stored at -80°C until use.

### Purification of mammalian CIV

Purification of bovine CIV was performed as described in (33) with some modifications (changes refer to the procedure described in section “Purification of cytochrome *c* oxidase” in ref. (33)). In the original protocol, separation of CIII and CIV involves centrifugation for 45 min at 100 000 × g. Here, the centrifugation time was 1 h at 126,000 × g. In the original protocol, after elution of CIV from the hydroxyapatite column and pooling of fractions containing the enzyme, the buffer is exchanged on a Sepharose CL-6B column. Here, this step was excluded and instead the pH was adjusted to 8.5 by titration with a 0.5 M Tris solution. In the next step, Triton X-100 was removed using Bio-Beads (Bio-Rad) instead of dialysis: Bio-Beads were added to the CIV fraction at 0.3 g/ml and stirred for 3 h at 4°C. The solution was then filtered to remove the Bio-Beads and the filtrate was centrifuged at 30,000 × g for 20 min at 4°C. The pellet was resuspended in 2% DDM, 20 mM Tris-HCl pH 8.5, 200 mM NaCl and centrifuged at 2000 × g for 20 min at 4°C. The supernatant containing CIV was then collected.

A HiTrap Q ion exchange column (GE Healthcare) pre-equilibrated with buffer (50 mM HEPES pH 7.4 and 0.08% [w/v] GDN) was used to exchange CIV into GDN using an Äkta Pure M25 chromatography system (GE Healthcare) operated at 4°C with UV detection at 280 nm and 420 nm. CIV was eluted with a gradient of increasing KCl (elution ∼400 mM KCl). Fractions containing CIV were concentrated with a 100-kDa molecular weight cutoff concentrator (Merck Millipore) and into 50 mM KCl, 50 mM HEPES pH 7.4 and 0.016% [w/v] GDN before being concentrated again. For cryo-EM samples, CIV was applied to a Superdex 200 Increase 10/300 GL size exclusion column to remove CIV aggregates. To make CHS solutions, 0.1% (w/v) CHS was mixed in 50 mM KCl, 50 mM HEPES pH 7.4, and 0.004% (w/v) LMNG or 50 mM KCl, 50 mM N-Cyclohexyl-2-aminoethanesulfonic acid (CHES) pH 9.0, and 0.004% (w/v) LMNG and sonicated for 4 min at 60% of the maximum power (Vibra-Cell VCX 130, Sonics & Materials) until it was clear, followed by centrifugation to remove large particles.

### Activity measurements

The activity of CIV was measured by monitoring the oxygen-reduction rate with an Oxygraph Clark-type oxygen electrode (Hansatech) operated at 25°C. A baseline oxygen concentration was first recorded with 50 μM oxidized cytochrome *c* from bovine heart, 10 mM ascorbate, 100 μM *N,N,N*’,*N*’-tetramethyl-phenylenediamine (TMPD) in 50 mM KCl, 50 mM HEPES pH 7.4, and either 0.016% (w/v) GDN or 0.05% (w/v) DDM. For the CHS titration, 0.004% (w/v) LMNG was used in the buffer instead of GDN or DDM with CHS concentrations in the range 165 μ M to 990 μ M and CIV was incubated in the buffer containing CHS prior to measurement. The reaction was initiated by addition of 15 nM CIV.

### Preparation of the two-electron reduced state and flash photolysis

CIV was diluted to 5 μM in 50 mM KCl, 50 mM CHES pH 9.0, and either 0.016% (w/v) GDN or 0.004% (w/v) LMNG with or without 0.06% (w/v) CHS. The samples were then transferred to a modified anaerobic Thunberg cuvette. The atmosphere was exchanged consecutively for N_2_ and CO on a vacuum line, after which the sample was incubated for 1–2 h. During that time, CO donates two electrons to each CIV, which results in formation of the “mixed-valence state” in which the catalytic site heme *a*_3_/Cu_B_ is reduced while heme *a* and Cu_A_ remain oxidized. Another CO molecule then binds to the reduced heme *a*_3_ resulting in formation of the CIV-CO complex (as monitored spectrophotometrically) (60): 2CO + H_2_O + Cu_B_^2+^Fe_*a*3_^3+^ --> CO_2_ + 2H^+^ + Cu_B_^+^ Fe_*a*3_^2+^-CO. The CO ligand was dissociated by a 10 ns laser flash (Nd-YAG laser at 532 nm, Quantel) and absorbance changes were monitored using a flash photolysis setup (Applied Photophysics, UK). The CO dissociation and recombination, and the following electron transfer from heme *a*_3_ to heme *a*, were monitored at 445 nm. The proton coupled electron transfer at a millisecond time scale was monitored at 598 nm, which is an isosbestic point for the CO-recombination reaction. The 598 nm traces were normalized to the absorbance changes associated with CO dissociation, measured at 445 nm.

### Preparation of the four-electron reduced CIV and flow flash

CIV was diluted to 5 μM in 50 mM KCl, 50 mM HEPES pH 8.0, and either 0.016% (w/v) GDN or 0.05% (w/v) DDM. The samples were transferred to Thunberg cuvettes, and the atmosphere was exchanged to nitrogen before reduction by 2 mM ascorbate and 1 μM of the electron mediator Phenazine methosulfate (PMS). After the samples were fully reduced, nitrogen was exchanged for CO. The flow-flash measurements were performed using a locally modified laser flash-photolysis system combined with a stopped-flow apparatus (39). Briefly, the reduced CO-bound enzyme was mixed at a ratio of 1:1 with an oxygen-saturated solution containing 50 mM KCl, 50 mM HEPES pH 8.0, and either 0.016% (w/v) GDN or 0.05% (w/v) DDM. The CO ligand was dissociated by a short laser flash (Nd-YAG laser at 532 nm, Quantel) at 0.2 s after mixing, which initiated the reaction. The reaction with oxygen was monitored at 445 nm.

### Grid preparation and cryo-electron microscopy

Purified CIV (35 μl) at a concentration of ∼8 mg/ml, was bubbled with carbon monoxide (Praxair) in a sealed container for ∼10 min to ensure an oxygen free environment. The sample was reduced by adding 3.5 μl of a mixture of 1.4 mM *N,N,N*’,*N*’-tetramethyl-*p*-phenylenediamine (TMPD, Millipore sigma) and 112 mM ascorbate (chem-impex international) then bubbled with carbon monoxide for another ∼10 min. The reduced CO-bound CIV was applied to homemade nanofabricated holey gold grids (61, 62) that had been glow-discharged in air (120 s, 20 mA using PELCO easiGlow). CryoEM specimen preparation was performed in a low light environment to prevent dissociation of carbon monoxide. Grids were blotted for 28 s at 4°C and 100 % humidity before rapid freezing in liquid ethane with a Vitrobot Mark IV (Thermo Fisher Scientific). Cryo-EM data were collected with a Titan Krios G3 electron microscope (Thermo Fisher Scientific) operated at 300 kV equipped with a prototype Falcon 4 direct detector device camera. Automated data collection was done with the EPU software package. An initial dataset of 4,222 movies, each consisting of 30 exposure fractions, was collected at a nominal magnification of 75,000×, corresponding to a calibrated pixel size of 1.03 Å. The camera exposure rate and the total exposure of the specimen were ∼4.7 e^-^/pixel/s and ∼42.7 e^-^ /Å^2^, respectively (***SI Appendix*, Table S1**).

### Cryo-EM image processing

All image analysis was performed with cryoSPARC v2 (63). Movies were aligned with *MotionCor2* (64) and contrast transfer function (CTF) parameters were estimated in patches with a 7×7 grid. The dataset was manually curated to remove movies with devitrification, large cracks in the ice, or poor CTF fit parameters, reducing the dataset size to 4,161 movies. Templates for particle selection were generated by 2D classification of manually selected particle images leading to 911,268 selected particles. After particle selection, images were corrected for local motion (65) and extracted in 256?256 pixel boxes (***SI Appendix* Table S1**). Extracted particle images were cleaned with four rounds of *ab initio* 3D classification and heterogeneous refinement, taking only the classes that corresponded to CIV after each round. This procedure further reduced the size of the dataset to 112,880 particle images, which were subject to non-uniform refinement to produce the final map. Chimera and coot were used to perform a ridged body fit of the CIV molecular model from a previous crystal structure (PDB ID:1V54) (49) and GDN into the final map.

### Classical molecular dynamics simulations

Classical atomistic molecular dynamics (MD) simulations were performed on the 13 subunit bovine CIV embedded in a lipid membrane comprising a 2:2:1 mixture of POPC, POPE, and cardiolipin constructed using CHARMM-GUI (66), The system was solvated in TIP3P water molecules, and 150 mM NaCl was added to neutralize the system. Water molecules in the D-pathway were modeled based on a prior x-ray structure (PDB ID:1V54). The total system comprised ∼250,000 atoms. To probe the effect of the sterol on the K-pathway dynamics, CIV was modeled with and without GDN, or with cholesterol at the GDN headgroup position seen by cryo-EM. The catalytic site was modeled in the O_H_ state (Fe^III^-OH^-^/Cu^II^-H_2_O Y_244_-O^-^), with heme *a* reduced or in the O_H,R_ state (Fe^III^-OH^-^/Cu^I^-H_2_O Y_244_-O^-^) with heme *a* oxidized. The protein, membrane, and water molecules were modeled using the CHARMM36 force field, whereas the redox cofactors were modeled using in-house parameters based on DFT calculations (67), with force field parameters (68) derived also for GDN. MD simulations were performed at *T*=310 K and *p*=1 atm with a 2 fs timestep, and treating long-range electrostatics with the particle mesh Ewald (PME) approach. All MD simulations were performed with NAMD2.14 and NAMD3.0. (69). See ***SI Appendix* Table S2** for the list of MD simulations.

## AUTHOR CONTRIBUTIONS

J.L.R. and P.B. designed research; J.D.T., A.M., D.R., P.S. performed research; J.D.T., A.M., D.R., P.S., V.R.I.K., J.L.R. and P.B. analyzed and interpreted the data; D.R, P.S. performed molecular simulations; J.L.R. supervised the electron cryomicroscopy studies; P.B. supervised the kinetic studies; V.R.I.K. supervised the computational studies; and J.D.T., J.L.R., P.B., V.R.I.K., A.M. wrote the paper.

## DECLARATION OF INTERESTS

### DATA DEPOSITION

Data deposition: The electron cryomicroscopy map described in this article has been deposited in the Electron Microscopy Data Bank (EMDB) (accession no. EMD-XXXX).

## ACKNOWLEDGEMENTS

This work was supported by the Knut and Wallenberg Foundation via grant 2019.0043 (PB) and Wallenberg Academy Fellow (WAF)-grant (VRIK), the Swedish Research Council grant 2018-04619 (PB), and the Canadian Institutes of Health Research grant PJT162186 (JLR). This work was also supported by the Swedish National Infrastructure for Computing (SNIC 2021/1-40) at Center for High Performance Computing (PDC) Centre, partially funded by the Swedish Research Council through Grant Agreement 016-07213. JDT was supported by a Canadian Institutes of Health Research Postdoctoral Fellowship and JLR was supported by the Canada Research Chairs program. Titan Krios cryo EM data were collected at the Toronto High-Resolution High-Throughput cryo-EM facility supported by the Canada Foundation for Innovation and Ontario Research Fund. Molecular graphics and analyses performed with UCSF Chimera, developed by the Resource for Biocomputing, Visualization, and Informatics at the University of California, San Francisco, with support from NIH P41-GM103311.

## REFERENCES

1. P. R. Rich, Mitochondrial cytochrome c oxidase: Catalysis, coupling and controversies. Biochem. Soc. Trans. 45, 813–829 (2017).

2. M. Wikström, V. Sharma, V. R. I. Kaila, J. P. Hosler, G. Hummer, New perspectives on proton pumping in cellular respiration. Chem. Rev. 115, 2196–2221 (2015).

3. A. A. Konstantinov, S. Siletsky, D. Mitchell, A. Kaulen, R. B. Gennis, The roles of the two proton input channels in cytochrome c oxidase from Rhodobacter sphaeroides probed by the effects of site-directed mutations on time-resolved electrogenic intraprotein proton transfer. Proc. Natl. Acad. Sci. U. S. A. 94, 9085–9090 (1997).

4. P. Brzezinski, P. Ädelroth, Pathways of proton transfer in cytochrome c oxidase. J. Bioenerg. Biomembr. 30, 99–107 (1998).

5. J. P. Hosler, et al., Polar residues in helix VIII of subunit I of cytochrome c oxidase influence the activity and the structure of the active site. Biochemistry 35, 10776–10783 (1996).

6. J. R. Fetter, et al., Possible proton relay pathways in cytochrome c oxidase. Proc. Natl. Acad. Sci. U. S. A. 92, 1604–1608 (1995).

7. R. B. Gennis, P. Brzezinski, Role of the Pathway through K (I-362) in Proton Transfer in Cytochrome c Oxidase. Biochemistry 2960, 2470–2476 (1998).

8. S. Yoshikawa, A. Shimada, Reaction Mechanism of Cytochrome c Oxidase. Chem. Rev. 115, 1936– 1989 (2015).

9. J. P. Hosler, S. Ferguson-Miller, D. A. Mills, Energy transduction: Proton transfer through the respiratory complexes. Annu. Rev. Biochem. 75, 165–187 (2006).

10. S. Ferguson-Miller, C. Hiser, J. Liu, Gating and regulation of the cytochrome c oxidase proton pump. Biochim. Biophys. Acta - Bioenerg. 1817, 489–494 (2012).

11. P. Brzezinski, A. Moe, P. Ädelroth, Structure and Mechanism of Respiratory III–IV Supercomplexes in Bioenergetic Membranes. Chem. Rev. 121, 9644–9673 (2021).

12. T. Tsukihara, et al., The whole structure of the 13-subunit oxidized cytochrome c oxidase at 2.8 A. Science 272, 1136–1144 (1996).

13. A. Maréchal, et al., A common coupling mechanism for A-type heme-copper oxidases from bacteria to mitochondria. Proc. Natl. Acad. Sci. U. S. A. 117, 9349–9355 (2020).

14. H. M. Lee, et al., Mutations in the putative H-channel in the cytochrome c oxidase from Rhodobacter sphaeroides show that this channel is not important for proton conduction but reveal modulation of the properties of heme a. Biochemistry 39, 2989–2996 (2000).

15. S. Arnold, The power of life-Cytochrome c oxidase takes center stage in metabolic control, cell signalling and survival. Mitochondrion 12, 46–56 (2012).

16. M. Hüttemann, I. Lee, L. I. Grossman, J. W. Doan, T. H. Sanderson, Phosphorylation of mammalian cytochrome c and cytochrome c oxidase in the regulation of cell destiny: respiration, apoptosis, and human disease. Adv. Exp. Med. Biol. 748, 237–264 (2012).

17. M. Hüttemann, B. Kadenbach, L. I. Grossman, Mammalian subunit IV isoforms of cytochrome c oxidase. Gene 267, 111–123 (2001).

18. L. Qin, C. Hiser, A. Mulichak, R. M. Garavito, S. Ferguson-Miller, Identification of conserved lipid/detergent-binding sites in a high-resolution structure of the membrane protein cytochrome c oxidase. Proc. Natl. Acad. Sci. U. S. A. 103, 16117–16122 (2006).

19. A. Musatov, Andrej, A. Robinson, Neal C., Bound Cardiolipin Is Essential for Cytochrome c Oxidase Proton Translocation. Biochimie 18, 1199–1216 (2014).

20. N. C. Robinson, L. H. Talbert, J. Zborowski, Cardiolipin-Depleted Bovine Heart Cytochrome c Oxidase: Binding Stoichiometry and Affinity for Cardiolipin Derivatives. Biochemistry 29, 8962– 8969 (1990).

21. C. Hiser, L. Buhrow, J. Liu, L. Kuhn, S. Ferguson-Miller, A conserved amphipathic ligand binding region influences K-path-dependent activity of cytochrome c oxidase. Biochemistry 52, 1385– 1396 (2013).

22. K. J. H. Van Buuren, B. F. Van Gelder, Biochemical and biophysical studies on cytochrome c oxidase. XIII. Effect of cholate on the enzymic activity. 333, 209–217 (1974).

23. I. Oleynikov, N. Azarkina, T. Vygodina, Direct Interaction of Mitochondrial Cytochrome c Oxidase with Thyroid Hormones: Evidence for Two Binding Sites (2022) https:/doi.org/10.20944/preprints202201.0328.v1.

24. C. Hiser, J. Liu, S. Ferguson-Miller, The K-path entrance in cytochrome c oxidase is defined by mutation of E101 and controlled by an adjacent ligand binding domain. Biochim. Biophys. Acta -Bioenerg. 1859, 725–733 (2018).

25. T. M. Ashton, W. Gillies McKenna, L. A. Kunz-Schughart, G. S. Higgins, Oxidative phosphorylation as an emerging target in cancer therapy. Clin. Cancer Res. 24, 2482–2490 (2018).

26. N. Simon, et al., Glucocorticoids decrease cytochrome c oxidase activity of isolated rat kidney mitochondria. FEBS Lett. 435, 25–28 (1998).

27. A. R. Vaz, et al., Bilirubin selectively inhibits cytochrome c oxidase activity and induces apoptosis in immature cortical neurons: Assessment of the protective effects of glycoursodeoxycholic acid. J. Neurochem. 112, 56–65 (2010).

28. P. S. Chae, et al., A new class of amphiphiles bearing rigid hydrophobic groups for solubilization and stabilization of membrane proteins. Chem. - A Eur. J. 18, 9485–9490 (2012).

29. S. Shimada, et al., Complex structure of cytochrome c –cytochrome c oxidase reveals a novel protein–protein interaction mode. EMBO J. 36, 291–300 (2017).

30. N. Yano, et al., X-ray structure of cyanide-bound bovine heart cytochrome c oxidase in the fully oxidized state at 2.0 Å resolution. Acta Crystallogr. Sect. FStructural Biol. Commun. 71, 726–730 (2015).

31. K. Shinzawa-Itoh, et al., Structures and physiological roles of 13 integral lipids of bovine heart cytochrome c oxidase. EMBO J. 26, 1713–1725 (2007).

32. L. Qin, D. A. Mills, L. Buhrow, C. Hiser, S. Ferguson-Miller, A Conserved Steroid Binding Site in Cytochrome c Oxidase. Biochemistry 47, 9931–9933 (2008).

33. U. Brandt, H. Schagger, G. von Jagow, Purification of cytochrome-c oxidase retaining its pulsed form. Eur. J. Biochem. 182, 705–711 (1989).

34. C. Hiser, J. Liu, S. Ferguson-Miller, BBA - Bioenergetics The K-path entrance in cytochrome coxidase is defined by mutation of E101 and controlled by an adjacent ligand binding domain. BBA - Bioenerg. 1859, 725–733 (2018).

35. H. Ilgü, et al., Variation of the detergent-binding capacity and phospholipid content of membrane proteins when purified in different detergents. Biophys. J. 106, 1660–1670 (2014).

36. J. Liu, C. Hiser, S. Ferguson-Miller, Role of conformational change and K-path ligands in controlling cytochrome c oxidase activity. Biochem. Soc. Trans. 45, 1087–1095 (2017).

37. P. Ädelroth, M. Svensson Ek, D. M. Mitchell, R. B. Gennis, P. Brzezinski, Glutamate 286 in Cytochrome aa3 from Rhodobacter sphaeroides Is Involved in Proton Uptake during the Reaction of the Fully-Reduced Enzyme with Dioxygen. Biochemistry 36, 13824–13829 (1997).

38. P. Ädelroth, P. Brzezinski, B.G. Malmström, Internal Electron Transfer in Cytochrome c Oxidase from Rhodobacter sphaeroides. Biochemistry, 2844–2849 (1995).

39. M. Brändén, et al., On the role of the K-proton transfer pathway in cytochrome c oxidase. Proc. Natl. Acad. Sci. U. S. A. 98, 5013–5018 (2001).

40. S. Hallén, P. Brzezinski, Light-induced structural changes in cytochrome c oxidase: implication for the mechanism of electron and proton gating. Biochim. Biophys. Acta 1184, 207–218 (1994).

41. S. Zong, et al., Structure of the intact 14-subunit human cytochrome c oxidase. Cell Res. 28, 1026–1034 (2018).

42. H. Aoyama, et al., The Whole Structure of the 13-Subunit Oxidized Cytochrome c Oxidase at 2.8 A. Science. 272, 1136–1144 (1995).

43. A. Punjani, D. J. Fleet, 3D variability analysis: Resolving continuous flexibility and discrete heterogeneity from single particle cryo-EM. J. Struct. Biol. 213, 107702 (2021).

44. I. Liko, et al., Dimer interface of Bovine cytochrome c oxidase is influenced by local posttranslational modifications and lipid binding. Proc. Natl. Acad. Sci. U. S. A. 113, 8230–8235 (2016).

45. R. Bolli, K. A. Nałecz, A. Azzi, The interconversion between monomeric and dimeric bovine heart cytochrome c oxidase. Biochimie 67, 119–128 (1985).

46. K. Shinzawa-Itoh, et al., Monomeric structure of an active form of bovine cytochrome c oxidase. Proc. Natl. Acad. Sci. U. S. A. 116, 19945–19951 (2019).

47. A. Musatov, N. C. Robinson, Cholate-Induced Dimerization of Detergent-or Phospholipid-Solubilized Bovine. Biochemistry 41, 4371–4376 (2002).

48. K. Shinzawa-Itoh, et al., Monomeric structure of an active form of bovine cytochrome c oxidase. Proc. Natl. Acad. Sci. 116, 19945–19951 (2019).

49. T. Tsukihara, et al., The low-spin heme of cytochrome c oxidase as the driving element of the proton-pumping process. Proc. Natl. Acad. Sci. U. S. A. 100, 15304–15309 (2003).

50. M. Svensson-Ek, et al., The X-ray crystal structures of wild-type and EQ(I-286) mutant cytochrome c oxidases from Rhodobacter sphaeroides. J. Mol. Biol. 321, 329–339 (2002).

51. J. Koepke, et al., High resolution crystal structure of Paracoccus denitrificans cytochrome c oxidase: New insights into the active site and the proton transfer pathways. Biochim. Biophys. Acta - Bioenerg. 1787, 635–645 (2009).

52. P. R. Rich, A. Maréchal, Functions of the hydrophilic channels in protonmotive cytochrome c oxidase. J. R. Soc. Interface 10 (2013).

53. M. Wikström, K. Krab, V. Sharma, Oxygen Activation and Energy Conservation by Cytochrome c Oxidase. Chem. Rev. 118, 2469–2490 (2018).

54. S. Supekar, V. R. I. Kaila, Dewetting transitions coupled to K-channel activation in cytochrome: C oxidase. Chem. Sci. 9, 6703–6710 (2018).

55. A. L. Woelke, A. Wagner, G. Galstyan, T. Meyer, E. W. Knapp, Proton transfer in the K-Channel analog of B-Type cytochrome c oxidase from Thermus thermophilus. Biophys. J. 107, 2177–2184 (2014).

56. I. Hofacker, K. Schulten, Oxygen and proton pathways in cytochrome c oxidase. Proteins Struct. Funct. Genet. 30, 100–107 (1998).

57. H. Lepp, E. Svahn, K. Faxén, P. Brzezinski, Charge transfer in the K proton pathway linked to electron transfer to the catalytic site in cytochrome c oxidase. Biochemistry 47, 4929–4935 (2008).

58. A. L. Woelke, G. Galstyan, E. W. Knapp, Lysine 362 in cytochrome c oxidase regulates opening of the K-channel via changes in pKA and conformation. Biochim. Biophys. Acta - Bioenerg. 1837, 1998–2003 (2014).

59. F. Kolbe, et al., Cryo-EM structures of intermediates suggest an alternative catalytic reaction cycle for cytochrome c oxidase. Nat. Commun. 12, 6903 (2021).

60. P. Brzezinski, B.G. Malmström, The reduction of cytochrome c oxidase by carbon monoxide. FEBS Lett. 187, 111–114 (1985).

61. C. R. Marr, S. Benlekbir, J. L. Rubinstein, Fabrication of carbon films with ∼500nm holes for cryo-EM with a direct detector device. J. Struct. Biol. 185, 42–47 (2014).

62. C. J. Russo, L. A. Passmore, Ultrastable gold substrates for electron cryomicroscopy. Science. 346, 1377 LP – 1380 (2014).

63. A. Punjani, J. L. Rubinstein, D. J. Fleet, M. A. Brubaker, CryoSPARC: Algorithms for rapid unsupervised cryo-EM structure determination. Nat. Methods 14, 290–296 (2017).

64. S. Q. Zheng, et al., MotionCor2: Anisotropic correction of beam-induced motion for improved cryo-electron microscopy. Nat. Methods 14, 331–332 (2017).

65. J. L. Rubinstein, M. A. Brubaker, Alignment of cryo-EM movies of individual particles by optimization of image translations. J. Struct. Biol. 192, 188–195 (2015).

66. S. Jo, T. Kim, V. G. Iyer, W. Im, CHARMM-GUI: a web-based graphical user interface for CHARMM. J. Comput. Chem. 29, 1859–1865 (2008).

67. M. P. Johansson, V. R. I. Kaila, L. Laakkonen, Charge parameterization of the metal centers in cytochrome c oxidase. J. Comput. Chem. 29, 753–767 (2008).

68. R. B. Best, et al., Optimization of the Additive CHARMM All-Atom Protein Force Field Targeting Improved Sampling of the Backbone ϕ, ψ and Side-Chain χ1 and χ2 Dihedral Angles. J. Chem. Theory Comput. 8, 3257–3273 (2012).

69. J. C. Phillips, et al., Scalable molecular dynamics on CPU and GPU architectures with NAMD. J. Chem. Phys. 153, 44130 (2020).

