## Supplemental figures for "Structural basis of mammalian Complex IV inhibition by steroids"

**This PDF file includes:**

Figure S1 to S3

Table S1 to S2

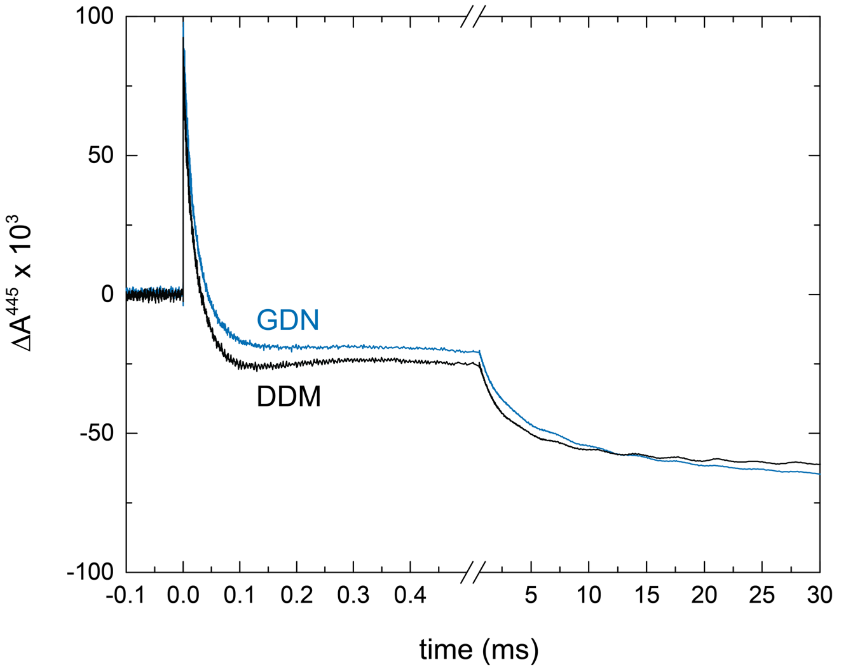


**Supplementary Figure 1**. Absorbance changes during reaction of reduced CIV with $O_2$. In this "flow-flash" experiment the four-electron reduced (anaerobic) CIV-CO complex is mixed with an $O_2$ solution. After ~30 ms a flash dissociates the CO ligand (at *t* = 0 in the figure), which allows $O_2$ to bind to heme *a*3 followed by step-wise oxidation of the CIV while $O_2$ is reduced to $H_2O$. The unresolved rapid increase in absorbance at *t* = 0 is associated with dissociation of the CO ligand, while the following kinetic components are associated with step-wise oxidation of the enzyme. The first component ($\tau \cong 10$ µs) is associated with binding of $O_2$ to heme $a_3$. It is followed by a further decrease in absorbance associated with formation of the so-called peroxy intermediate ($\tau \cong 50$ µs). The slight increase in absorbance in the range 0.1-0.3 ms is associated with fractional electron transfer from $Cu_A$ to heme *a* with a time constant of ~100 µs during the peroxy to ferryl transition at the catalytic site. The final, slow decrease in absorbance with a time constant of ~2 ms is associated with formation of the fully oxidized CIV. The last two reaction steps are linked to proton uptake through the D-pathway. Oxidation of the reduced CIV is not associated with proton uptake through the K-pathway, but formation of the peroxy state requires charge compensation through this pathway (1) Therefore, the slowed kinetic component associated with formation of this intermediate in the GDN sample is indicative of an impaired K-pathway. The data show that CIV is fully oxidized over the same time scales in GDN and DDM, which indicate that neither intramolecular electron transfer nor proton transfer through the D-pathway is slowed in GDN. The mixing ratio of the $O_2$-saturated buffer to reduced enzyme was 1:1. The time delay between mixing and laser flash was 200 ms. Both the reduced CIV sample and $O_2$ buffer contained 50 mM KCl, 50 mM HEPES pH 8.0, and either 0.016% GDN or 0.05% DDM. A laser artefact has been removed for clarity (large apparent negative absorbance due to the laser hitting the detector at *t* = 0).

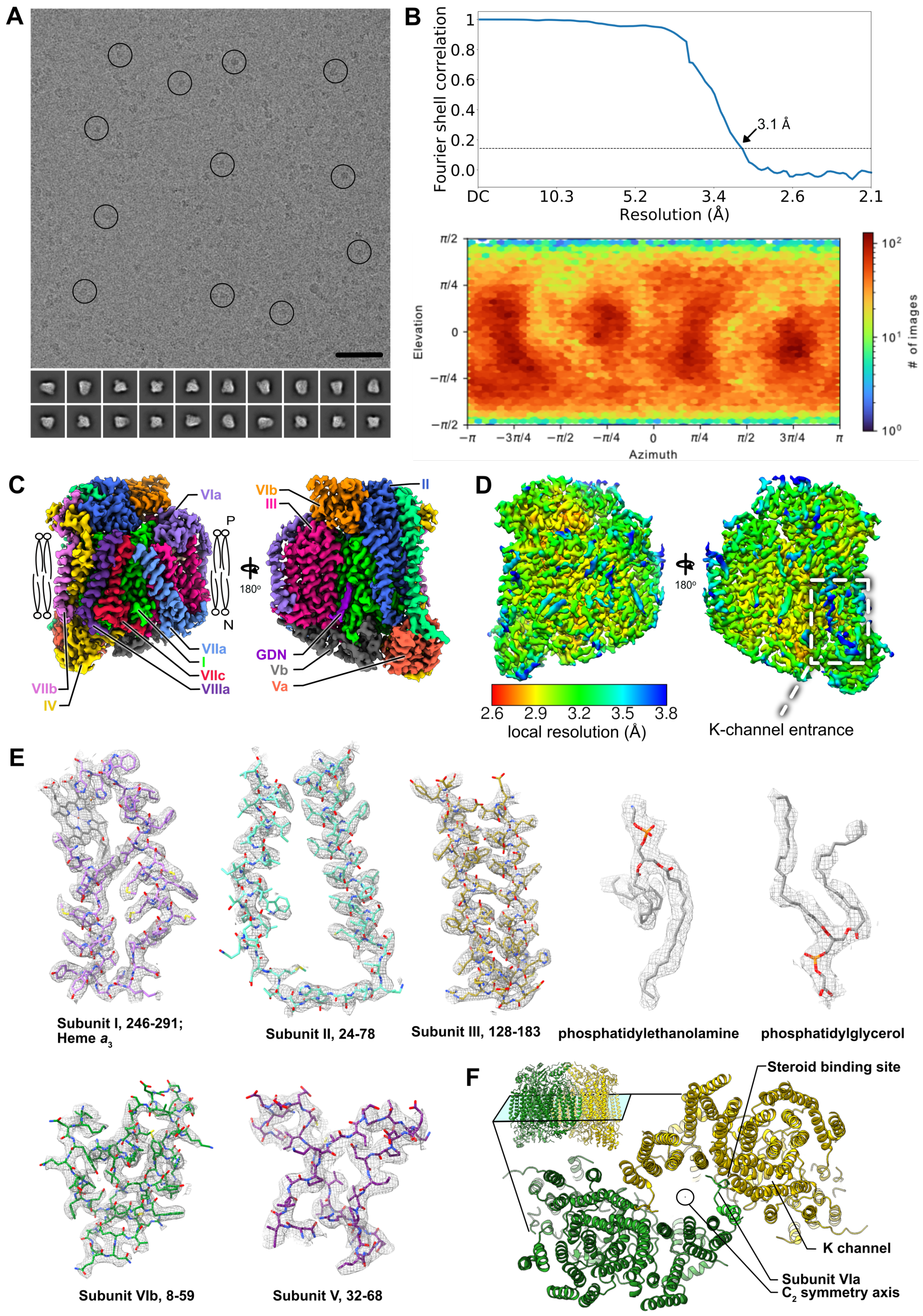


**Supplementary Figure 2**. (**A**) Representative micrograph and 2D class average images for CIII₂. Scale bar, 500 Å. (**B**) Fourier shell correlation (FSC) curve after correction for solvent masking (*upper*) and viewing direction distribution for particle images (*lower*). (**C**) Cryo-EM map of CIV with all subunits labelled. (**D**) Local resolution in the CIV map. (**E**) Examples of model in map fit. (**F**) $CIV_2$ dimer from a previous crystal structure (2) showing K-pathway and N-terminal tail of coxVIa occupying steroidal binding site.

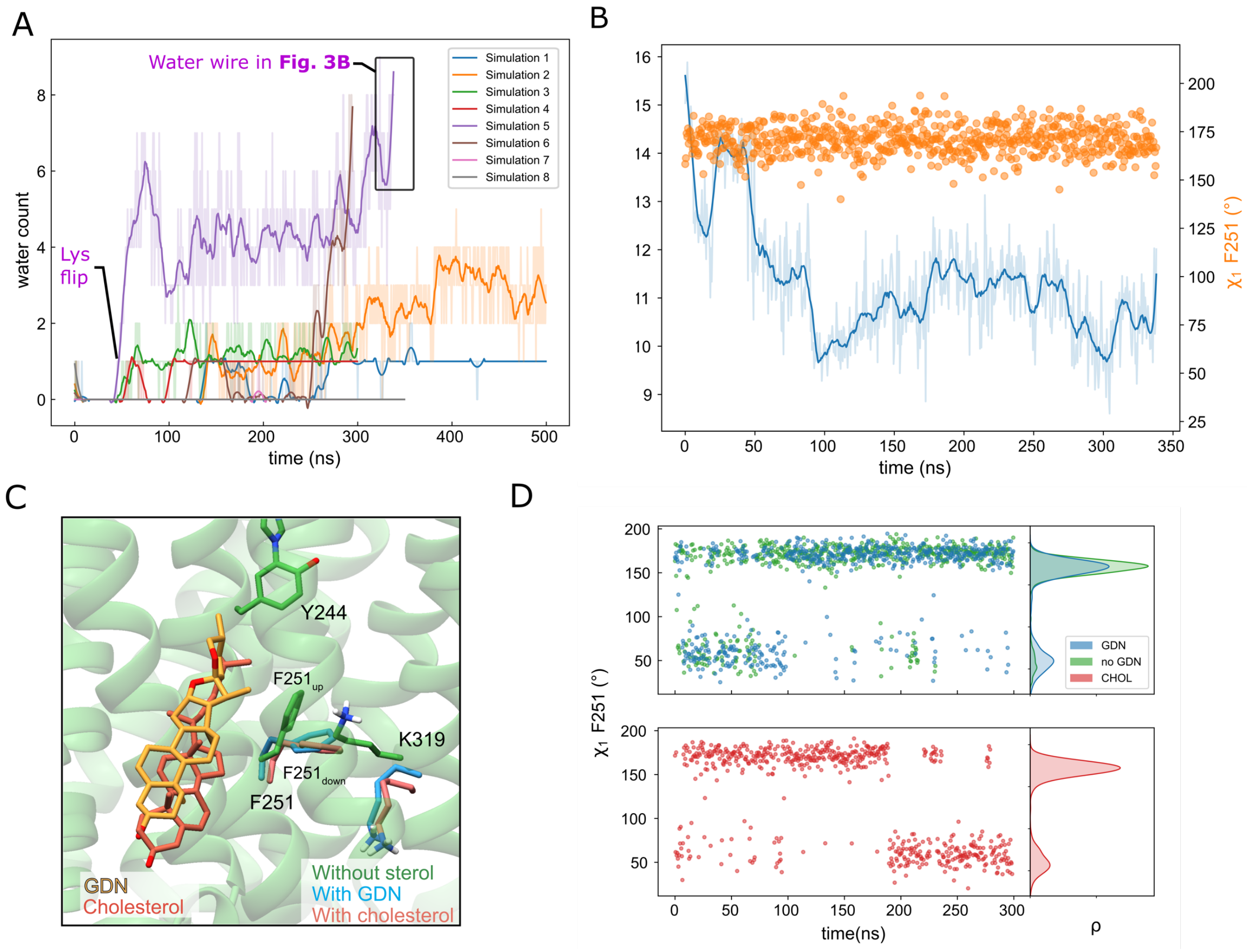


**Supplementary Figure 4**. (**A**) Hydration of the K-pathway in different MD trajectories (see SI Appendix, Table S2). Transition of Lys319 to the 'up' conformation in simulation 5 (without bound sterol, see SI Appendix, Table S2) is coupled to rapid hydration of the K-pathway. Water molecules within 5 Å of Thr316 were counted. (**B**) Phe251 dihedral and Lys319-Tyr244 distance during simulation 5. Phe251 samples only the up conformation ($\chi_1$=175°) when Lys319 points towards the active site. (**C**) GDN/cholesterol binding site next to the K-pathway showing Phe251 and Lys319 conformations in simulations with GDN (in blue), with cholesterol (in red), and without a bound sterol (in green). (**D**) Distribution of the Phe251 dihedral angle ($\chi_1$) in simulations with GDN (GDN), cholesterol (CHOL), and without sterol (no GDN). The down conformation of Phe251 ($\chi_1$=60°) is significantly enhanced with the bound sterol.

**Supplementary Table 1.** Cryo-EM data acquisition and image processing.

| **Data Collection** | |
|---|---|
| Electron Microscope | Titan Krios |
| Camera | Falcon 3 |
| Voltage (kV) | 300 |
| Nominal Magnification | 75,000 |
| Calibrated physical pixel size (Å) | 1.03 |
| Total exposure (e/Å$^2$) | 42.7 |
| Exposure rate (e/pixel/s) | 4.7 |
| Number of frames | 30 |
| Defocus range (µm) | 0.9 to 2 |
| **Image Processing** | |
| Motion correction software | *MotionCor2* |
| CTF estimation software | *cryoSPARC v2* |
| Particle selection software | *cryoSPARC v2* |
| Micrographs used | 4,161 |
| Particle images selected | 112,880 |
| 3D map classification and refinement software | *cryoSPARC v2* |

**Supplementary Table S2**. List of performed molecular dynamics simulations. The catalytic site was modelled in the $O_H$ state ($Fe^{III}$-$OH^-$/$Cu^{II}$-$H_2O$ $Y_{244}$-$O^-$ with heme *a* reduced) and the $O_{H,R}$ state ($Fe^{III}$-$OH^-$/$Cu^{I}$-$H_2O$ $Y_{244}$-$O^-$ with heme *a* reduced).

| **Name** | **Redox state** | **Sterol** | **Length (ns)** | **Comment** |
|---|---|---|---|---|
| S1 | $O_H$ | - | 500 | - |
| S2 | $O_H$ | GDN | 500 | - |
| S3 | $O_{H,R}$ | - | 300 | - |
| S4 | $O_{H,R}$ | GDN | 300 | - |
| S5 | $O_{H,R}$ | - | 350 | $Glu62_B$ protonated |
| S6 | $O_{H,R}$ | GDN | 300 | $Glu62_B$ protonated |
| S7 | $O_{H,R}$ | cholesterol | 300 | - |
| S8 | $O_{H,R}$ | cholesterol | 350 | $Glu62_B$ protonated |

### References


1. M. Brändén, *et al.*, On the role of the K-proton transfer pathway in cytochrome c oxidase. *Proc. Natl. Acad. Sci. U. S. A.* **98**, 5013–5018 (2001).
2. N. Yano, *et al.*, X-ray structure of cyanide-bound bovine heart cytochrome c oxidase in the fully oxidized state at 2.0 Å resolution. *Acta Crystallogr. Sect. FStructural Biol. Commun.* **71**, 726–730 (2015).
